# Robust Optical Flow Algorithm for General, Label-free Cell Segmentation

**DOI:** 10.1101/2020.10.26.355958

**Authors:** Michael C. Robitaille, Jeff M. Byers, Joseph A. Christodoulides, Marc P. Raphael

## Abstract

Cell segmentation is crucial to the field of cell biology, as the accurate extraction of cell morphology, migration, and ultimately behavior from time-lapse live cell imagery are of paramount importance to elucidate and understand basic cellular processes. Here, we introduce a novel segmentation approach centered around optical flow and show that it achieves robust segmentation by validating it on multiple cell types, phenotypes, optical modalities, and in-vitro environments without the need of labels. By leveraging cell movement in time-lapse imagery as a means to distinguish cells from their background and augmenting the output with machine vision operations, our algorithm reduces the number of adjustable parameters needed for optimization to two. The code is packaged within a MATLAB executable file, offering an accessible means for general cell segmentation typically unavailable in most cell biology laboratories.

## INTRODUCTION

It is hard to state how important live-cell microscopy has been for our understanding of biology. Currently, researchers are afforded a humbling window into the sheer complexity of cell behavior thanks to advancements in both the quality and diversity of optical modalities available for live-cell microscopy^1^. Great strides have been made on the front-end to obtain both higher quality data and larger amounts of it. However, the field has quickly realized a serious issue on the back-end – viable methods to actually analyze the imagery in meaningful and robust ways are often lacking^2,3^. In terms of live-cell imaging, this analysis often means segmenting the cell through time lapse imagery, with the “holy grail” being robust segmentation applicable to both labeled and label-free cells to control for perturbations to the cells observed^4–6^. Indeed, the symptoms of this problem are readily apparent in the field wherein biologists are often so focused on the difficult task of ensuring that their experimental design and execution is accurate, that often times little bandwidth is left over for analyzing the exquisite data strenuously collected. As a result, many complex biological functions are often over simplified – the fluid-like motions of the cell membrane during migration are treated like a rigid body by tracking the cell *via* a nucleus, or the dynamic process of cell adhesion is measured by the projected cell area at a single time point are common examples.

Such data is not left on the table willingly, but rather done so because it is a *hard* problem. This problem has garnered an immense amount of interest towards solving^7^, which, in turn ironically causes its own problem: researchers already overwhelmed with the complexity of the data they have painstakingly collected are oftentimes inundated by the sheer number of purported solutions to their problems. In our own experience, many hours can be invested before realizing a well-intentioned published segmentation method is simply inadequate when applied to your own data set. It is perhaps unsurprising then that cell segmentation *via* a laborious manual tracking approach is still considered the gold standard.

Great advances have indeed been made in cell segmentation^7,8^, however, many methods follow an apparent common trend – they perform well only when applied to the type of imagery they were designed for^2^. Machine learning has recently become an appealing avenue for a more general method of cell segmentation^4,9–11^. These methods require initial supervision through manually segmented cells to teach the algorithms to correctly classify cells later on in an unsupervised matter. However, training can be time consuming, and it remains an open question as to how robust the library must be in order to achieve robust segmentation capabilities^12,13^.

A viable solution for robust segmentation is to exploit aspects that are present in all live-cell microscopy, regardless of the type of optical modality and subsequent generated imagery: cellular motion and dynamic morphology. Indeed, nearly all current methods utilize the image intensity (*I*) as a function of position (x, y) within in a single image, disregarding information as to how this intensity function varies with time (*t*) and re-evaluating each image as if they are completely unrelated to the previous or next. This strikes us as a rather incomplete strategy in terms of robustness – when one considers the sheer diversity of live-cell imagery, the one unifying characteristic is that live-cell microscopy is dynamic with respect to time.

Here, we show that by leveraging the fact that live cells move and are morphologically dynamic, optical flow-based segmentation algorithms can offer quite robust and simple means to segment live-cell imagery. Optical flow-based methods have been previously employed to quantify live-cell imagery, largely in the context of movement of tagged proteins within a given cell^14–16^. We are aware of exceptionally few cases in which optical flow is utilized for label-free segmentmentation^17^. To this end, we introduce a label-free cell segmentation technique based on the Färneback method^18^, and show that leveraging cellular movement via optical flow is an appealing strategy that accurately segments cells across cell types, phenotypes, optical modalities, resolution and environments with relatively few parameters required to optimize. This robust algorithm is computationally inexpensive, simple to use, and packaged within MATLAB, offering accessibility into dynamic cell behavior oftentimes unavailable to typical cell biology labs

## RESULTS

The foundation of the label-free segmentation algorithm outlined here is optical flow, in which the spatial changes in image intensity *I* (x, y) between two consecutive time frames is quantified to characterize relative movement. The underlying principle is that at every pixel a displacement is estimated that maps the image at time frame *t* to the image at time frame *t*+Δ*t*. This assumes that while pixels can move in x and y, their net intensity *I*(x, y) remains constant, or changes negligibly between consecutive frames. More formally this can be written out as:

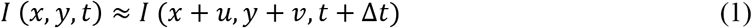

Where *u* and *v* are the local displacements of the local image regions at *x* and *y* after time Δt. Another way to think of it is the pixels in image *t* are pushed to form the pixels in image *t*+Δ*t* by the optical flow field. Applying a first-order Taylor expansion to the right hand side of (1) and dividing by Δ*t* yields^19^:

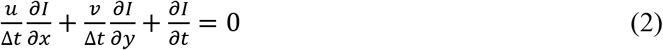

In which 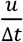 and 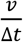 are the optical flow in *x* and *y*, respectively. Equation (2) is known as the optical flow constraint equation, and the flow field calculated from it is then used as a means to estimate which pixels correspond to a cell in a given image: pixels in which there is no movement between consecutive frames do not generate much displacement and can be considered background, while pixels in which larger displacement are generated can be considered a cell, given that certain criteria are met. This approach does not depend on any particular image characteristic in *I*(*x, y*), but rather how *I*(*x, y*) changes between consecutive frames, and thus enables highly robust cell segmentation without the need for labels.

Optical flow is a general term for a variety of strategies that try to measure relative intensity displacement between consecutive images, and methods are optimized to compute either sparse or dense flow fields for either small or large object displacements. For instance, rigid objects that do not undergo shape changes between consecutive frames (i.e. cars captured on traffic cameras) are well suited for methods which employ sparse flow fields, such as Lucas-Kanade^20^. For imagery of deformable objects that may change shape between consecutive frames (for instance, a cell with filopodia extending or retracting), it is necessary to employ a dense flow field method that estimates flow within objects as well as around the perimeter. Similarly, objects that may make relatively large displacements between consecutive frames are best dealt with methods that utilize pyramidal multi-scale resolution techniques. Considering that the movement of cells can vary significantly between cell type or experimental set up and lead to relatively large cell displacements between consecutive frames, we have found the Farnebäck method to work exceptionally well for estimating flow fields of live-cell imagery.

A detailed description of the Farnebäck method is given elsewhere^18^, but the key elements are 1) the use of multi-level resolution pyramid and 2) the use of quadratic polynomials to quantify the intensity of pixel neighborhoods. Figure 1A depicts this resolution pyramid, in which a grouping of four pixels are averaged to form a single pixel at the next resolution level, with the higher pyramid levels corresponding to lower resolutions of the imagery. At the lowest resolution level, neighborhoods of pixels (i.e. 5×5 square) are fitted with a quadratic polynomial weighted towards the center of the neighborhood for each time frame. It is assumed that a single displacement can describe the transformation of the fitted polynomial neighborhood at frame *t* to the fitted polynomial neighborhood at frame *t* + Δ*t*, and this displacement is iteratively calculated. The final displacement is a 2D vector that describes the flow of intensity in one neighborhood between consecutive images, and is then used as *a priori* knowledge for the fitting of quadratic polynomials at the next pyramid level below it (higher resolution). The displacement is then recalculated based on polynomial fits of newer, higher resolution neighborhoods, and in turn is used as *a priori* knowledge for the pyramid level below it, and so on. The end result is a 2D vector field that characterizes the optical flow at the true resolution of the image that is quite resilient to noise, deformable objects, and large displacements. Once the flow field is calculated at the highest resolution, a threshold is then applied to the magnitude, shown in Figure 1B. All pixels corresponding to a flow value below the threshold are considered background, while pixels with a flow value above the threshold are segmented as cells. The remaining pixels are those that exhibit significant optical flow and represent moving cells, and are subsequently grouped together and filled to create a binary mask (Figure 1C) that can then be used to segment cells from their background (Figure 1D).

**Figure 1:**
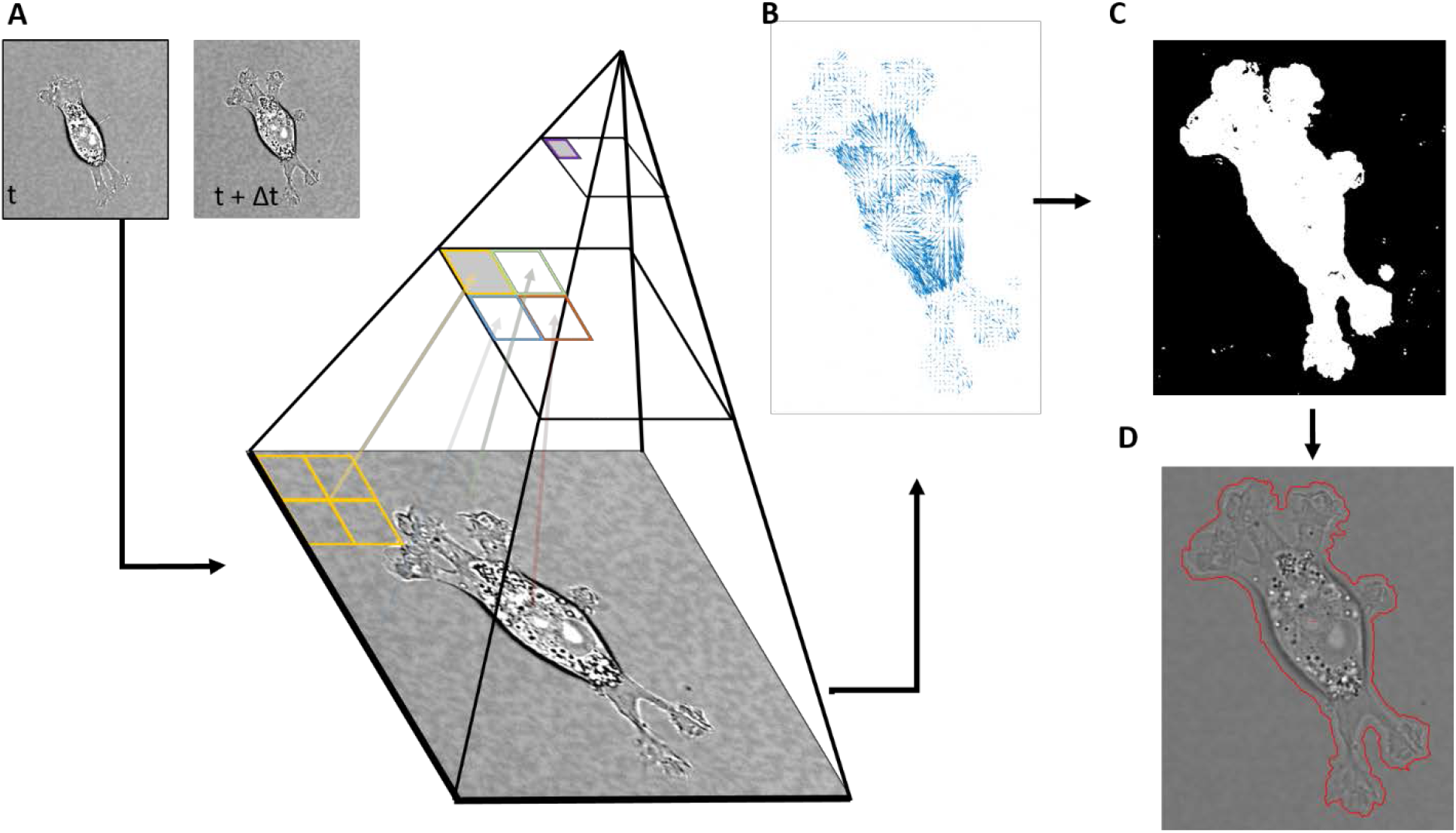
Overview of the segmentation algorithm. A) Two consecutive image frames are used to calculate the optical flow via the Farnebäck algorithm, which utilizes a multi-level resolution pyramid to estimate intensity displacements between the two frames. B) Once the flow field is calculated, a threshold Th is applied to the magnitude of the individual flow vectors, leaving only flow vectors with a magnitude above Th. C) The pixels associated with the remaining flow field form a binary mask to estimate where the cell is in the image. D) The mask is closed and filled to create the segmented perimeter of the cell.

The Optical Flow algorithm has a single primary parameter that requires optimization for label-free cell segmentation: the optical flow vector magnitude threshold (Th). Additionally, two secondary computer vision parameters, a size filter, and to a lesser extend smoothing disk can be altered depending on the imagery. Each parameter is intuitive and takes little time to optimize for a given data set. High Th values only segment areas in which large changes in pixel intensity due to object motion (i.e. flow), and too high of Th values can lead to an underestimation of cell area (Figure 2A). Similarly, too low of Th values can pick up smaller intensity changes outside of a cell due to stochastic fluctuations in cameras or experimental artifacts and can often times lead to an over estimation of cell area. The size filter simply prohibits the segmentation of objects smaller than the entered value to eliminate the segmentation of cellular debris, precipitates, or other objects common *in-vitro*. Finally, the smoothing disk is a standard morphological operation for filling small holes and smoothing the segmented perimeter of the cell (Figure 2B). Oftentimes, the size filter and smoothing disk are robust with regards to cell types or optical modalities, leaving only the threshold (Th) to optimize. For a given experimental set up/cell type, a single Th value is often robust enough to capture a variety of different cell morphologies within a single field of view, as is shown in Figure 2C.

**Figure 2:**
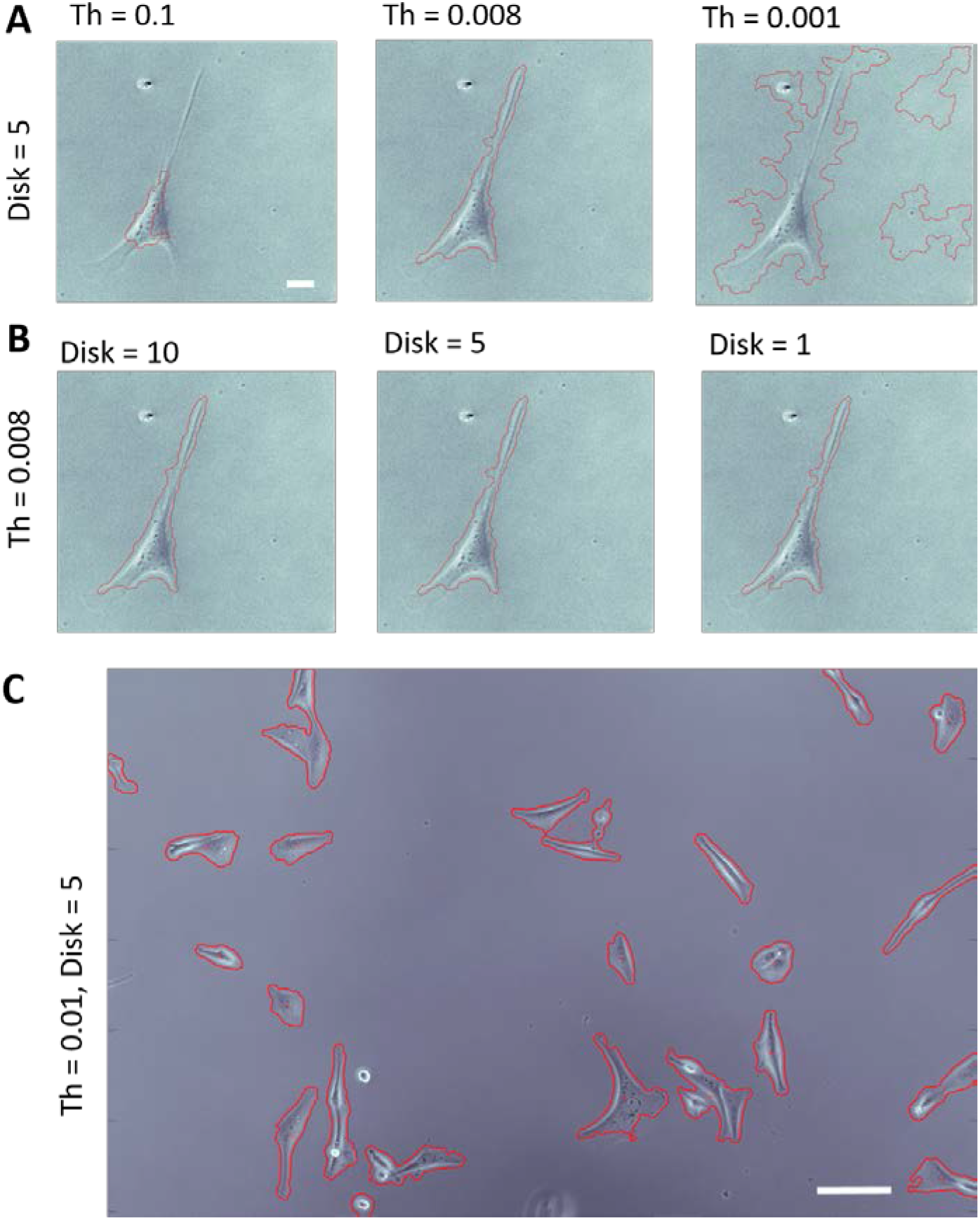
The effect of optical flow parameters on segmentation of a single Hs27 fibroblast under 10x phase contrast microscopy. A) The optical flow threshold, Th, can lead to over- or under-segmentation if not optimized, and is typically the only parameter that needs to be optimized for a given data set. Scale bar 20μm B) The smoothing disk of the segmented cell effects the perimeter, but often times does not drastically alter the accuracy of the segmentation itself. C) A larger field of view of MBA-MD-231 cells under 10x phase showing a single value of Th = 0.01 can adequately segment cells under a variety of morphologies. Scale bar 50μm.

To validate the optical flow segmentation algorithm and demonstrate its robustness, segmentation of a variety of cell types and optical modalities are shown in Figure 3. To serve as a baseline for typical segmentation techniques, a fluorescently labeled A549 cell is easily segmented based on the cell’s motion between consecutive time frames rather than the fluorescent tag in Figure 3A. Next, Dictyostelium cells are segmented without the use of any labels under transmitted light (TL) microscopy that is standard for most cell biology laboratories in Figure 3B. Dictyostelium was chosen to highlight the ability of optical flow to segment many different morphologies, behaviors, or otherwise phenotypes any given cell type exhibit, as cells of different phenotypes can exhibit drastically different morphologies and thus image characteristics^21,22^. In Figure 3B, Dictyostelium cells exhibiting amoeboid, keratocyte/fan, or oscillator/intermediate phenotypes are segmented, each of which has a distinct morphology and migratory characteristics. Amoeboid phenotypes typically possess a rounded morphology leading to high contrast boundaries around their edge when imaged under TL or phase contrast microscopy, whereas keratocyte phenotypes exhibit a flatter, broad lamellipodia that typically have much lower contrast at the cell boundary comparatively. The intermediate phenotype switches between morphologies that look similar to both amoeboid and keratocyte, and all three phenotypes are segmented accurately via optical flow despite these differences in morphology and image characteristics.

**Figure 3:**
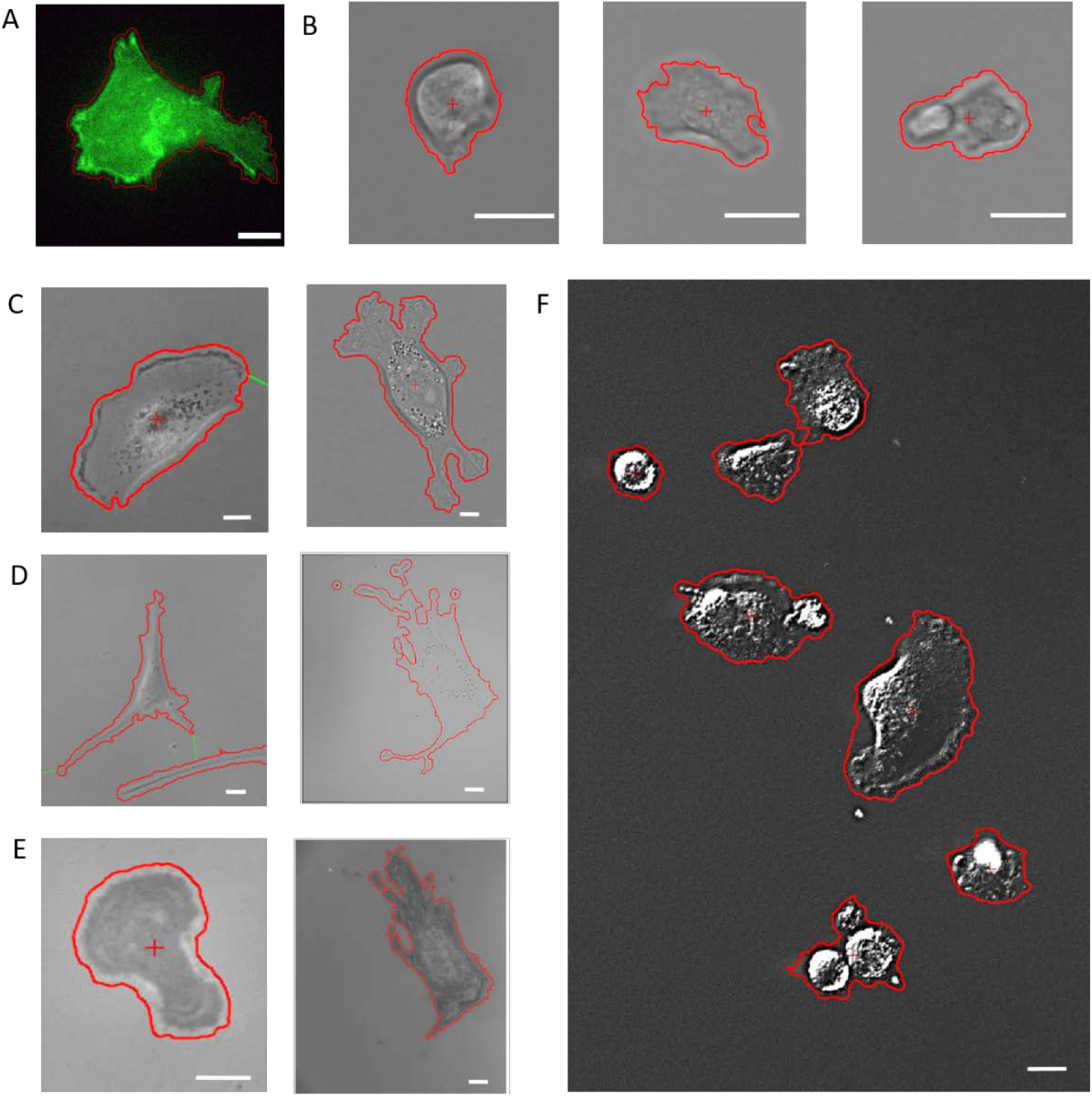
Robust segmentation via optical flow across optical modalities, cell types, and phenotypes. Scale bars are 10μm. A) A549 carcinoma cell stably transfected with green fluorescence to serve as a baseline, 40x. B) Dictyostelium cells under 40x TL exhibited a range of phenotypes: Amoeboid (left), keratocyte (center), and intermediate (right). C) MDA-MB-231 cells accurately segmented under 10x phase contrast (left) or 40x TL (right). D) Hs27 Fibroblasts accurately segmented under 10x phase (left) or 40x TL (right). E) Both MDA-MB-231 cell (left) and Hs27 fibroblast (right) segmented under IRM. F) Multiple MDA-MB-231 cells featuring a variety of morphologies accurately segmented under a single threshold value at 20x DIC. All images have been contrast enhanced for better visualization.

Similarly, fibroblasts (Hs27) and epithelial (MDA-MB-231) cells were segmented under TL, phase contrast, Differential Interference Contrast (DIC), and Interference Reflection Microscopy (IRM). Phase, DIC, and TL are commonly used modalities in most cell biology labs, with cells imaged under TL having much lower contrast compared to phase or DIC typically. IRM is a lesser employed, but powerful modality. Figure 3C & 3D show both cell types under phase and TL modes, covering a range of complex morphologies between the two cell types. The optical flow algorithm is able to segment all cells tested regardless of morphology or optical modality due to the use of cell motion between consecutive images as a means of segmentation. Figure 3C shows two different typical morphologies of MDA-MB-231 cells exhibit – a spread out morphology in phase contrast and a more complicated, five lamellipodia morphology under transmitted light that are both accurately segmented via optical flow. Similarly, the optical flow algorithm is able to segment fibroblasts exhibiting multiple lamellipodia in complex geometries accurately under both phase and TL in Figure 3D. Even within the same field of view, the optical flow algorithm is able to segment cells of different morphology, as shown in Figure 3F where multiple MDA-MB-231 cells are segmented under 20x DIC, spanning clumped, balled up, spread and fan-like morphologies with the same flow threshold.

IRM is a unique modality that visualizes interference patterns generated from reflections of an incident beam of light as it passes through materials of different refractive indices, highlighting the interactions at the cell-substrate interface^23^. In Figure 3E, both MDA-MB-231 and Hs27 cells are segmented accurately via optical flow despite having extremely different intensity or imagery characteristics compared to more traditional modes of microscopy. Indeed, IRM can reveal intricate cellular structures oftentimes not readily visible under transmitted light^24^, and it is worthwhile to note that the multi-modality capability of the optical flow algorithm allows for direct comparison of observed cell behavior in the case of IRM vs TL. Figure 4 shows a direct comparison of the segmentation via optical flow of a fibroblast under both TL and IRM shown in Figure 3E, highlighting the different aspects of the fibroblast each modality is suited to detect. Such a comparison could be useful considering that it is common for cell adhesion to be characterized by measuring the area of spread cells, however it has been shown that the area observed under IRM is directly related to the degree and strength of cell adhesion^24^.

**Figure 4:**
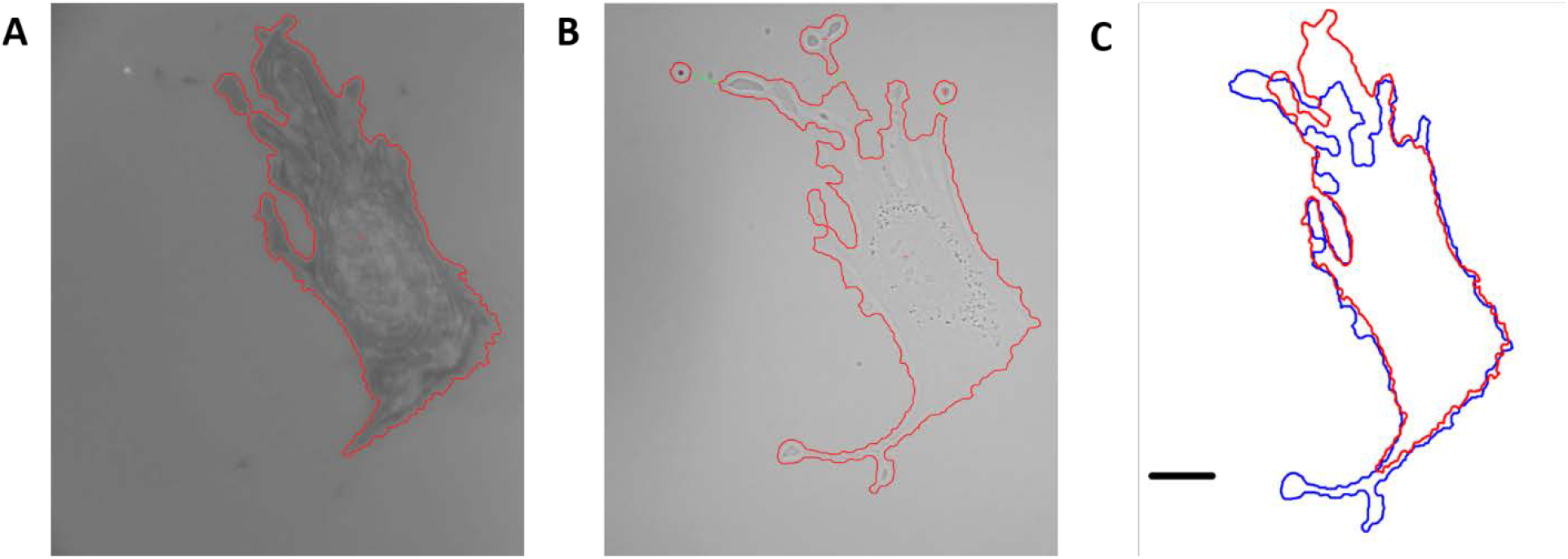
Direct comparison of a Hs27 fibroblast under A) 40x IRM and B) 40x TL. IRM highlights the cell-substrate interface while TL projects the entire 3 dimensional cell onto a 2 dimensional image, leading to fairly different segmented areas, compared in C). Scale bar is 20um.

A unique capability arises as a consequence of utilizing cell motion as a means for segmentation - the ability to distinguish cells in complex, yet stationary environmental surroundings. This is of particular interest as it is becoming increasingly common for research groups to observe cells interacting with various fabricated surfaces and structures as a means to elucidate underlying mechanisms governing cell behavior. In-vitro platforms investigate fundamental cellular processes such as adhesion and migration through a plethora of two or three-dimensional surfaces/structures including printed lines of ligands^25^, nanodots^26^, nanopillars^27^, and 3D matrices^28^. Each one of these techniques results in a unique architecture present in images that pose a distinct challenge for segmenting cells interacting with these structures, often times resulting in the use of fluorescent or manual labeling to distinguish cellular boundaries from fabricated structures. One such example are platforms that investigate contact guidance, in which repeating grooves are etched or fabricated into a substrate to create three dimensional structures to investigate cellular response to topographical cues^29,30^.

Our group recently introduced a contact guidance platform capable of integrating with nearly all forms of live-cell microscopy^31^, which generates a unique problem for accurate cell segmentation against the backdrop of etched grooves that serve as topographical cues. Figure 5 shows Hs27 fibroblasts atop such grooves under IRM, in which the etched grooves can scatter/diffract light differently depending on their topographical dimensions, leading to imagery much more complex than a traditional flat substrate or petri dish. The issue is further complicated by the fact that the relative intensity/pixel values of the topographical structures can be quite similar to those of a cell, making accurate segmentation exceptionally challenging. However, due to the stochastic fluctuations of cell motion, our optical flow algorithm segments the cell with reasonable accuracy against the backdrop of the etched structures with a threshold value of Th = 0.01-0.02. To the best of our knowledge, the presented optical flow algorithm is the only option for label-free segmentation of cells interacting with such structures in a highly generalizable manner.

**Figure 5:**
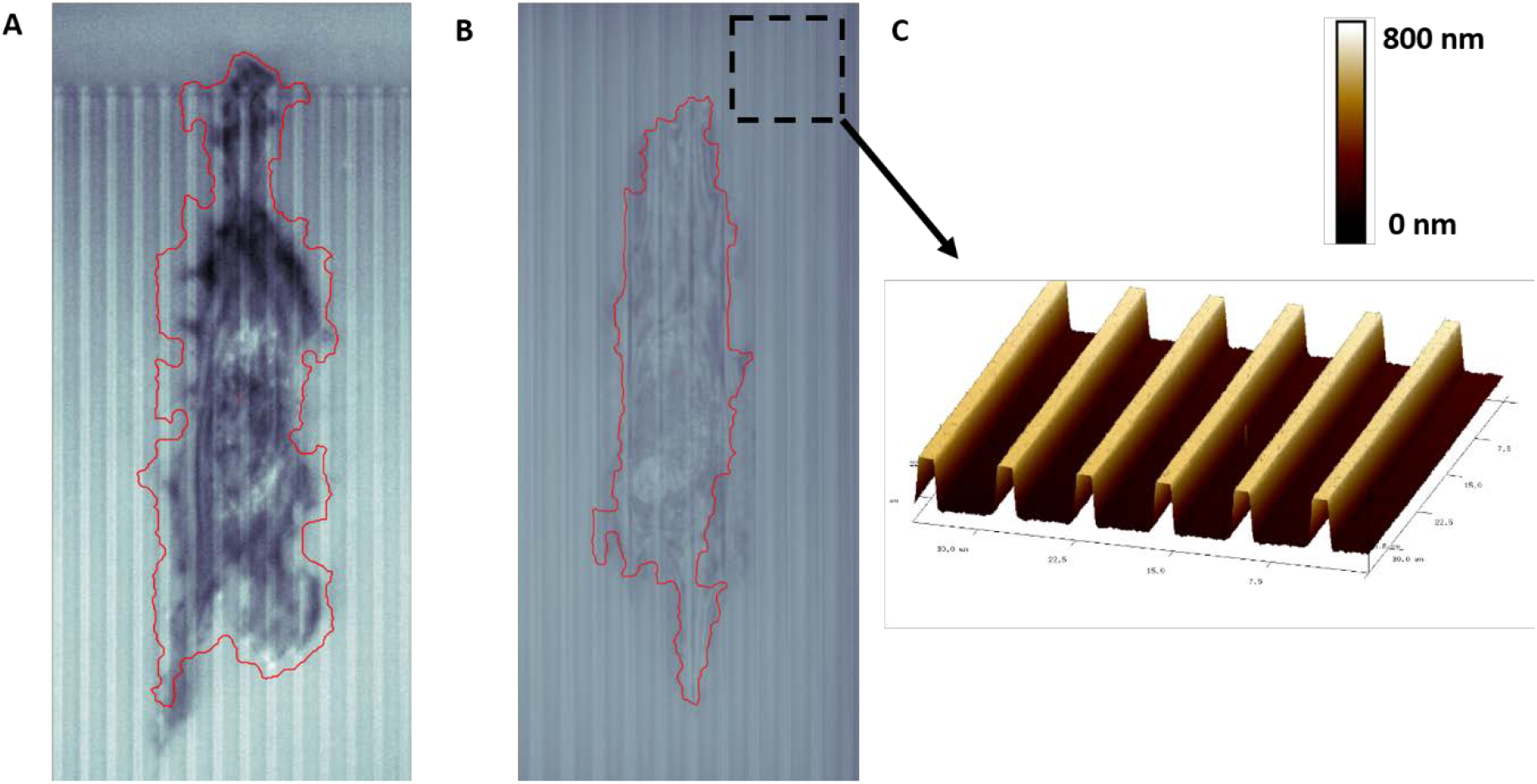
Hs27 under IRM with a 40x objective, atop etched contact guidance grooves of depth (D), ridge (R) and groove (G) dimensions A) D = 0.330 um, R= 2 um, G = 4 um, and B) D = 0.725 um, R = 2 um, G = 8 um. A flow threshold of Th = 0.01 (A) or 0.02 (B) is able to segment the fibroblast with reasonable accuracy under diverse in-vitro imagery. C) Atomic Force Microscope image depicting three-dimensional view of the contact guidance grooves.

## DISCUSSION

Here we have introduced an optical flow based strategy for label-free segmentation of cells. By utilizing the changes in image intensity (*I*) as a function of position (x, y) and time (t) as a means to differentiate areas that belong to cells versus the background, the burden of segmentation is largely removed from cell contrast and shifted towards cell movement. This is a novel way of interpreting live-cell imagery, and one can think of this as additional information that exists for every pixel in a typical image histogram, but has been seemingly overlooked. Thus, accurate segmentation is readily achieved without any specificity or otherwise training with regards to optical modality, cell type, cell behavior, or in-vitro environment/structures, creating a truly robust strategy for cell segmentation. The use of the term cell “movement” is perhaps misleading, as it infers a drastic (i.e. noticeable) translation of the cell position between time frames. However, in our experience, even cells that appear stationary between time frames exhibit significant optical flow within the boundary of their area due to intensity fluctuations from intracellular processes that are readily segmented, usually with the same flow threshold Th suitable for segmentation of motile cells within the same experimental/optical/imagery conditions. These benefits combined with relatively few and simple parameters that require tuning make optical flow segmentation strategies appealing to the broader cell biology community.

Optical flow based strategies outlined here do however have some shortcomings and limitations. First, the accurate segmentation under optical flow requires an extremely stable experimental set up such that the only movement between consecutive frames is that of the cells and not the experimental background itself. As such, drift of the microscope stage or objective focus can lead to a large displacement of pixel intensity values, and thus large values of optical flow that are unrelated to actual cell motion. Image alignment software can be employed to take care of drift in lateral movements (x, y) of the stage, however, drifts in focus (z) are extremely detrimental to optical flow based segmentation. On top of the experimental set up, the data acquisition or camera set up also plays a crucial role in determining the quality of optical flow segmentation. The illumination intensity and exposure time should be optimized to produce an intensity histogram that is not over or under exposed – optical flow simply cannot be calculated accurately from pixels which are truncated at maximum or minimum values. Along similar lines, the frequency of data collection is an important factor that can directly affect the quality of segmentation under optical flow. Two of the underlying assumptions of the optical flow algorithm is that the net intensity is conserved and that the net displacement is small between consecutive frames. Thus, the longer the time between frames, the more objects (i.e. cells) move and the less likely these assumptions are to be valid, and ultimately the quality of the segmentation suffers. In our experience with the variety of cells/experiments outlined in this manuscript, time steps of 10 min yielded similarly accurate cell segmentation as time steps of 20s, however this is likely to be highly dependent upon the cell type and experimental conditions.

Another drawback, common to label-free segmentation approaches, is there is no inherent way to distinguish if a segmented object is a single cell or a multicellular aggregate (i.e. Figure 3F). This makes it necessary to incorporate a manual selection process to separate single from multi-cell aggregates when dealing with segmenting many cells or precaution when setting up an experiment to seed cells at a low enough density such that cell-cell contact is negligible. Finally, by utilizing changes in intensity as a means of segmentation, the level of accuracy of the segmentation will depend on optical artifacts that can change or shift prevalence depending on optical modality, cell, and even phenotype. For instance, under phase-contrast microscopy, and in particular at low magnification (i.e. 10x) objectives, the well-known halo artifact in which cell boundaries are obscured by high contrast intensity changes can lead to an overestimation of segmented area depending on the cell morphology. Future work could address optical modality specific artifacts to improve the efficacy of the outlined algorithm under such conditions (i.e. phase contrast halo correction).

## CONCLUSION

The algorithm presented here, to the best of our knowledge, is the first optical-flow based label-free technique that offers relatively simple and robust means of cell segmentation. The notion of utilizing cell motion as a means to distinguish cells from their background is a rather elegant strategy for segmentation in a variety of environments or optical modalities, without the need for labels. It is worth noting that the use of optical-flow based segmentation is not exclusive to other image processing techniques, opening the possibility of optical-flow to be combined with segmentation techniques, such as machine learning, for potential improvements in segmentation accuracy, robustness and ultimately classification. With robust segmentation capabilities and few parameters to optimize, the ease-of-use of this optical flow segmentation algorithm stands to offer accessibility into the dynamic behavior of cells for typical cell biology laboratories across disciplines.

## MATERIALS & METHODS

### Cell cultures

All human cell lines were purchased through ATCC (Hs27 #CRL-1634, MDA-MB-231 #HTB-26, A549 #CCL-185), and cultured according to ATCC protocols in DMEM (ATCC #30-2002) in 10% fetal bovine serum (ATCC #30-2020) at 37°C in 5% CO_2_. Cells were subcultured according to ATCC protocols and cells were harvested between 30-80% confluence for all experiments. Wild-type *Dictyostelium discoideum* cells of the AX2 strain generously obtained from the Devreotes laboratory (Johns Hopkins University, USA) were used in this study and were cultured axenically in HL5 media at 22°C as outlined in^21^. For experiments involving *Dictyostelium* imaging, cells were harvested at ~80% confluence by gently aspirating/rinsing the culture dish/flask and using the supernatant of suspended cells for live cell imaging.

### Microscope/in vitro set ups

Live cell imaging was performed using phase contrast (10X, 0.9 NA objective), Differential Interference Contrast (DIC, 20X, 0.8 NA objective transmitted light (TL 40X, 1.4 NA objective), or interference reflection microscopy (IRM) (40X, 1.4 NA objective). For mammalian cell lines: a heated stage and temperature controlled enclosure held the stage temperature at 37.0 ± 0.04°C (Zeiss) with humidity and CO2 regulated at 98% and 5%, respectively, by flowing a gas-air mixture through a heated water bottle and into the enclosure. For *Dictyostelium* imaging, cells were imaged in glass-bottomed petri dishes at room temperature (22°C). Focus was stabilized for the multi-hour long experiments using an integrated hardware-based focus correction device (Zeiss Definite Focus). All mammalian cell line experiments were done in serum free media, and conducted on either glass-bottomed well plates/petri dishes or in the case of contact guidance structures, quartz chips detailed previously^31^. The *Dictyostelium* imaging was conducted on glass bottomed petri dishes in HL5 culture media.

## ACKNOWLEDGEMENTS

The authors gratefully acknowledge the Devreotes laboratory of Johns Hopkins University for the *Dictyostelim discoideum* cell line. M.C.R. gratefully acknowledges support from the National Research Council Research Associateship Program and the Jerome and Isabella Karle Distinguished Scholar Fellowship Program. Funding for this project was provided by the Office of Naval Research through the Naval Research Laboratory’s Basic Research Program and by the Biological Technology Office of the Defense Advanced Research Program Agency.

